# Rabies prevention and control practice and associated factors among dog owners in Aksum town and Laelay-Machew district, north Ethiopia: community based comparative cross-sectional study

**DOI:** 10.1101/436493

**Authors:** Letebrhan Gebrezgiher, Gebretsadik Berhe, Aregawi Gebreyesus Belay, Alefech Adisu

## Abstract

**Background:** Rabies is nearly 100% fatal zoonotic disease. One thousand seven hundred suspected rabies exposures reported in north Ethiopia, Tigray region in 2017, which has the highest rabies prevalence from Ethiopia. Almost half of them were from Central zone only. Of these 38% were in Aksum town and Laelay-Machew. Though Rabies exposure is prevalent in Tigray, there is scanty information on rabies prevention and control practices among dog owners. Thus, this deals with rabies prevention and control practice and associated factors among dog owners.

**Methods and materials:** Comparative community based cross-sectional study was conducted in Aksum town and Laelay-Machew district from March 01 to 20, 2018. A multi stage sampling was employed to recruit 558 households. Data were collected via structured and pretested questionnaire. Data were entered into Epi_info_7 and then exported to SPSS_20 for analysis. Both descriptive and inferential analysis was done with 95% confidence intervals at p value of 5% for the final model.

**Result:** The overall prevalence of poor rabies prevention and control practice was 56% [95%CI (50, 61.9)] in urban and 62% [95% CI (57.2, 67.7)] in rural dog owners. In urban; being government employee [AOR (95%CI) = 0.35 (0.13, 0.94)], private employee [AOR (95%CI) =0.39(0.16, 0.97)] and having poor attitude [(AOR (95%CI) =1.84 (1.04, 3.25)] were significantly associated with outcome variable. Whereas in rural dwellers; having no formal education [AOR (95%CI)=6.41(1.1,38.6)], poor attitude [AOR (95%CI)= 2.19 (1.18,4.05)], having one dog [AOR (95%CI)=3.31(1.34,8.15)], travel ≥30 minute to get vaccine [AOR (95%CI)= 4.26 (2.14,8.47]), no history of dog bite exposure [AOR (95%CI)= 4.16(1.49,11.6)] and neighbors as their source of information [AOR (95%CI) =3.64 (1.31,10.1)] have statistical significance with the outcome variable.

**Conclusion and Recommendation:** The prevalence of poor rabies prevention and control practice was higher among rural dog owners. Thus; interventions should be implemented both to urban and rural residents based on the identified findings so as to promote effective rabies prevention and control activities.

## INTRODUCTION

Rabies is a zoonotic disease caused by virus in the Rhabidovirus family of the genus Lyssavirus. It transmits to human through close contact with saliva (bite or scratch) of infected animals[1, 2]. Carnivorous such as dogs, cats, foxes, jackals, bats, raccoons and skunks are rabies reservoirs depending on the continents. But, Dogs are responsible for more than 95% of all rabies transmission to humans in developing countries [3].

Rabies disease is widely distributed across continents of the world. Globally, It causes around 60,000 human deaths per year despite more than 15 million people receive post exposure prophylaxis (PEP). Above 95% of deaths occur in Asia and Africa. Africa accounts for 44% of deaths. More than 40% of deaths occur in children under 15 years old [4, 5]. It was assumed that more than 2,700 human lives lose estimated annually in Ethiopia in 2015 [5]. By the year 2010- 2013, About 4,734 dogs had vaccinated, 3,550 ownerless dogs killed by chemical applicants, 388 dogs sterilized and for 7,050 students awareness creation programs performed in Mekelle city and its surrounding [6]. Yet, rabies is still prevalent in Ethiopia particularly in Tigray region. There were 2,928 cases exposed to suspected rabid dog and 31 rabies deaths in Ethiopia. Of these 1,439 (49.1%) were in Tigray region ranked first from the other regions as the national surveillance data reported by Ethiopian public health institute in 2016 [7]. As the regional health bureau annual report, about 1,723 persons exposed to rabies suspected dog were reported in Tigray region. The highest proportion of cases 827 (48%) were reported from central zone. Of which individuals exposed to suspected rabid dogs were highly reported in Aksum town 314 (38%) and but relatively lower 25 (0.03%) in Laelay Machew district in 2016/2017 [8].

Although rabies is 100% preventable but, nearly 100% fatal disease once its clinical symptoms appear. Globally, it leads to over 3.7 million disability-adjusted life years and an estimated of 8.6 billion dollars economic loss annually due to premature death (55%) and direct cost of PEP (20%) [5, 9].

Rabies elimination is feasible through dog vaccination and prevention of dog bites. Dog vaccination is the preferred method of controlling and eliminating rabies worldwide. According to world health organization (WHO), dog vaccination coverage should at least 70% in rabies endemic zones to eradicate/block outbreak occurrence [2]. In most African countries, where dog vaccination is not free of charge, the coverage is as low as 9% (Tanzania) [10]. Anti-rabies vaccines are expensive and consequently out of reach for many peoples in developing countries[11]. However; recent successes of rabies control through mass vaccination of dogs have been reported in Philippines, Bangladesh and South Africa [12].

Dogs were the reason for 97% of rabies related human deaths in Ethiopia. Despite the availability of vaccines both to human and dogs in the country, only 3.9% dogs were found vaccinated as the study done from 2009-2012 in Addis Ababa [13]. The widespread use of traditional medications and religious approach to treat rabies cases as evil spirit are also the challenges for prevention and control of rabies. Rabies victim individuals especially from rural areas usually come to health institutions after failing traditional intervention and loss of life from family members [14].

Prevention of human rabies needs community effort involving both veterinary and public health officials. Baseline information on the coverage rates of practices (dog vaccination, dog restrain, [15] proper care of bite wounds, seeking PEP and management of rabid dogs and carcasses) is important to predict rabies control and elimination efforts of the urban and rural dwellers [9].

There is a scarcity of information on rabies prevention and control practice and associated factors among dog owners in Ethiopia, Knowledge on rabies, formal employment, higher education, male headed households, urban residence, history of dog bite and vaccine availability were identified as significant factors for rabies prevention and control practices in the studies conducted in Tanzania, Kenya, Hawasa, Addis Ababa and Bahirdar [13, 16–18]. Knowledge, attitude and practice (KAP) towards rabies prevention and control found to be associated with sex, age, level of education and occupation of study participants in Debretabor town and Debark district [19, 20].

Descriptive studies conducted in Gonder Zuria, Gonder town and Dessie city assessed the overall KAP towards rabies. But, these didn’t identify the contributing factors to rabies prevention and control practice [21–23]. A cross sectional studies conducted in Hawasa town, Jimma zone, Debretabor, Bahirdar and Debark are tried to assess the significant factors for KAP on rabies. But these are not well informative in terms of the strength, precision and direction of the association between variables and did not compare among urban and rural households [18–20, 24, 25]. In most studies even though people were familiar with rabies, there is still a gap in practices towards rabies prevention and control measures. Therefore, this study was aimed to assess rabies prevention and control practice and associated factors among dog owners in urban (Aksum town) and rural (Laelay-Machew district).

## Methods and Materials

### Study setting and design

This study was conducted in Aksum town and Laelay-Machew district, Tigray region, Ethiopia from March 01 to 20/2018. Aksum town is located in Central Zone of Tigray Regional State, Northern Ethiopia 248 km from Mekelle the regional capital. The town has five Kebelles with a total population of 74,007 out of which 34,376 are males and 39,631 females as obtained from Woreda health office profile. Aksum town has two health centers, one general, one referral hospital and one veterinary clinic. Laelay-Machew district is a rural district found in the surrounding of Axum town having 16 kebeles with a total population of 80,445 of these 41,027 are females and 39, 418 males. The district has five health centers and 16 health posts with 16 biannual outreach sites for dog vaccination. There was no available data of dog population and vaccination coverage rates in both Woredas.

Community based comparative cross sectional study was employed to compare Rabies prevention and control practice status and associated factors.

### Study population

All dog owner households in randomly selected ketenas of Aksum town and Laelay-Machew district during the study period.

Household heads or their spouses of ≥18 years age (in the absence of the household head) who lived at least for six months as permanent resident in the study area and having dog at least three months old prior to this study were included in the study. However, Dog owners’ household head or their spouses aged ≥18 years old who have communication problem were excluded from the study.

### Sample size and procedure

Sample size calculation was determined based on the risk factors of practice on rabies prevention in different literatures. A double proportion formula was used to calculate sample size by applying Epi info stat calc version 7. The following assumptions were considered in calculating sample size: 95% confidence interval, 80% power, 1:1 ratio of exposed to non-exposed group, OR of 2.7 and taking the proportion of P1=81.1% of urban households and P2=92.1% of rural households and multiplied by 1.5 of design effect and added 10% of non-response rate and then became 558 participants [17] (Table 1).

Multistage sampling technique was employed for the selection of study unit. The study areas were selected based on burden of suspected rabies exposures. There were five kebeles in Aksum town and 16 kebeles in Laelay Machew district. At 3^rd^ stage, data was collected using systematic random technique from the selected ketenas/ Kushet every third for rural and second household for urban after randomly select the first household (figure 3).

**Table 1:**
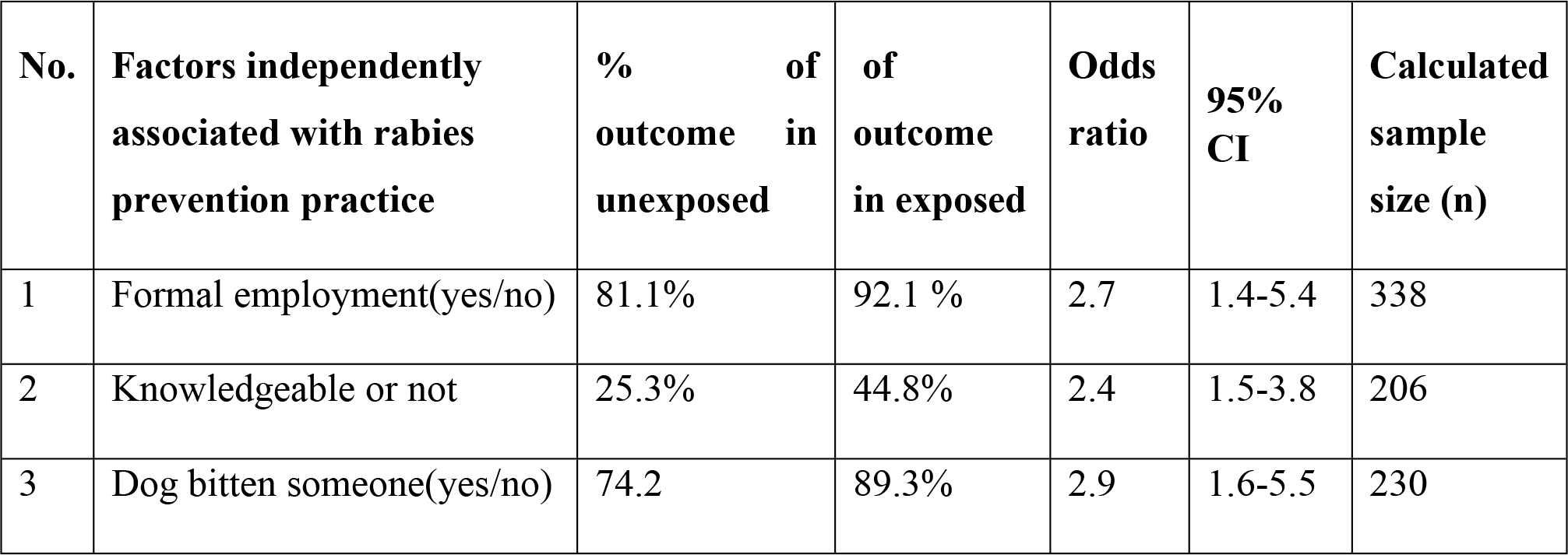
Sample size determination by EPI-Info for the second objective (factors associated with rabies prevention and control practice)

### Data collection tool and technique

A validated tool was adopted from previous studies [17, 18, 20, 28]. Data were collected using a structured and pretested questionnaire by trained interviewer. Primary data on socio-demographic characteristics of households, six knowledge questions on rabies (description of the disease, mode of transmission, outcome, range of species affected and means of prevention and control), ten attitude questions and six practice questions towards rabies prevention and control strategies (dog vaccination, dog restrain, timely seeking PEP, first aid, action for suspected rabid dog and carcasses management) collected from the selected households. The questionnaire was developed first in English version and translated to local language (Tigrigna) for local appropriateness and easiness in approaching the study participants and it was also back translated to English by third person to check the consistency. Household heads or their spouses were interviewed on all over the questionnaire content. Data was collected by seven trained health extension workers and two supervisors (health officers).

### Data quality control

Two day training was given to the data collectors on such issues as the techniques of data collection (face to face interview skills) and supervisors on how to check the completeness and consistencies of the questionnaires filled by the data collectors to ensure data quality. The questionnaire was pre-tested on 5% (28) of the sample size out of the sampled ketenas both in the urban and rural study areas then the questions that induce ambiguity based on the response obtained was revised and rephrased. Moreover; skip pattern and sequencing of questions was incorporated based on the pretest finding. The principal investigator visited and made close follow up to the data collectors to ensure whether appropriately collecting the data to prevent bias and give day to day feedback. At the end of each day filled questionnaire validity and completeness was checked before they return back from field. The principal investigator also verified the data completeness before data entry and analysis stages. Data was cross checked using double data entry.

### Analysis

After collecting the questionnaire, data was cleaned and checked for its completeness. After complete checkup the data coded and entered in to Epi-info version 7 software’s up on creating the questionnaire template. The data was exported to Stastical package for social sciences (SPSS) version 20 software for analysis. The frequency distributions of both dependent and independent variables were done using descriptive statistics (frequencies, mean, standard deviation and percentage) and presented in the form of text and tables. Cross tabulation was made using Pearson chi square test to select variables entered to binary logistic regression. Odds ratio (OR) and confidence interval at 95% was computed to test the association between independent and dependent variables. Variables with p-value of ≤ 0.25 cut of point in bivariate analysis were entered to logistic model for multivariable analysis. Normality was checked for continuous variables using histograms. Multicollinearity was checked using variance inflation factor (VIF) and no variables with VIF >10. Goodness of fit was examined using the Hosmer-Lemeshow goodness-of-fit test, at P-value >0.05 was accepted. Three separate multivariable models fitted. P-value <0.05 were considered statically significant relationship for identifying independent significant factors.

### Variables and measurements

**Dependent variable;** Rabies prevention and control practice (Good/poor)

**Independent variables; Socio-demographic/economic variables** such as; Sex, age, educational status, occupation, marital status, religion, ethnicity, residence (urban/rural), monthly income, household size and number of dogs in the household was taken.

**Health service related factors** such as; Distance from veterinary clinic, and availability of service

**Personal factors** such as previous history of dog bite to self and any family members, knowledge (good/poor), attitude towards rabies prevention and control (good/poor) and source of information.

## RESULT

A total of 558 dog owner households (279 in Aksum town and 279 in Laelay Machew district) participated in the study making a response rate of 100%.Of these The overall prevalence of poor rabies prevention and control practice was 56% [95%CI (50, 61.9)] in urban and 62% [95% CI (57.2, 67.7)] in rural dog owners. The overall mean score of practice was 8.7 (±3.5) with range of (0-16) in urban respondents and 9.0 (±3) with the range of (3-17) in rural ones.

### Characteristics of respondents: Socio demographic, Personal and health service, and factors on practice towards rabies prevention and control

The mean age of respondents Aksum town was 44.7 (±13.4 standard deviation) and Laelay Machew was 46.9 (±11.8SD). About 174 (62.4%) respondents in Aksum town and 242 (86.7%) in Laelay Machew district were married.

Of the total respondents 96 (34.4%) in Aksum town were private employee and 257 (92.1%) respondents in Laelay Machew district were farmers. About 127 (45.5%) of respondents in Aksum town and 113 (40.5%) of respondents in Laelay Machew district had no formal education. About 273 (97.8%) respondents in Aksum town and 278 (99.6%) in Laelay Machew district were Orthodox Christian followers (Table 2).

**Table 2:**
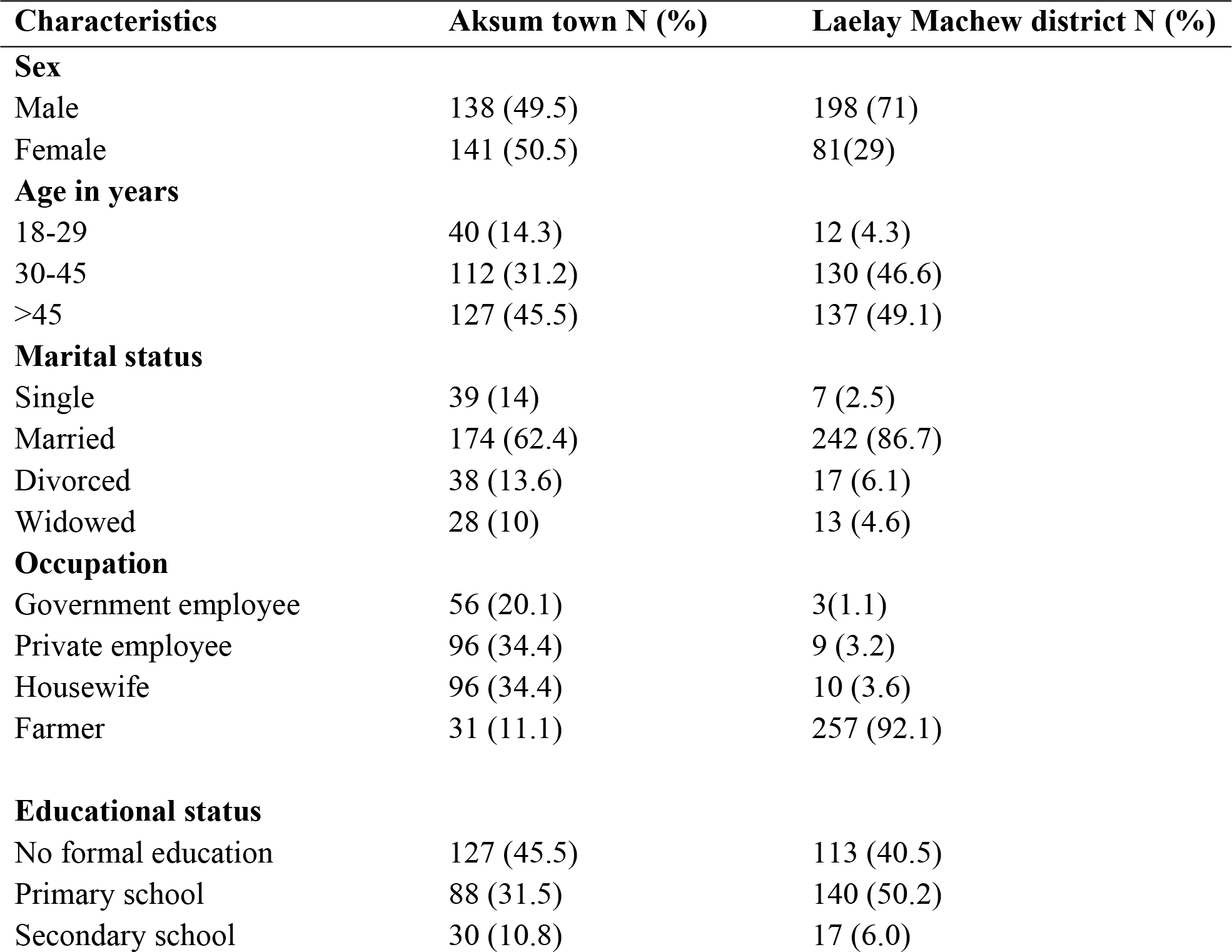
Socio demographic characteristics of respondents in Aksum town and Laelay Machew district, Central zone, Tigray region, Ethiopia, March 01-20/2018

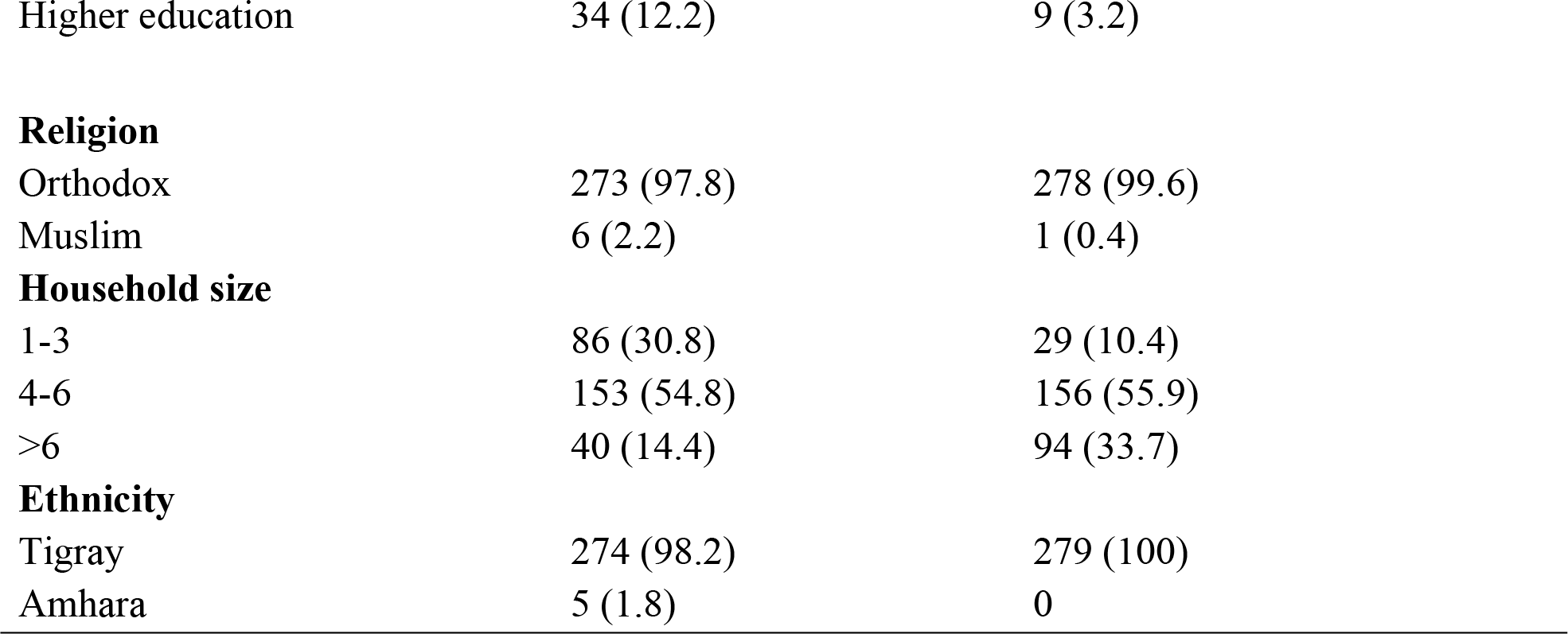

About two hundred eleven (75.6%) of urban and 119 (42.7%) of rural respondents, their source of information was health professional about rabies. The overall mean score of respondents knowledge on rabies disease was 8.7 (±1.1) in urban and 8.3 (±1.3) in rural respectively. More than half 177 (63.4%) of urban and 148 (53%) of rural respondents were above the mean score level as having good knowledge on rabies prevention and control.

The overall mean attitude score was 7.7 (±1.3) in urban and7.4 (±1.3) in rural. About 178 (64%) of urban and 131 (47%) of rural respondents had good attitude towards rabies prevention and control by scoring above the mean attitude score level. Out of the total participants, 45 (16.1%) of urban and 21 (7.5%) rural respondents were ever bitten by a suspected rabid dog. About 52 (18.6) of urban respondents and 25 (9%) rural respondents had family members ever bitten by dog. One hundred six (38%) rural respondents travel ≥30 minutes to get dog vaccine (Table 3).

**Table 3:**
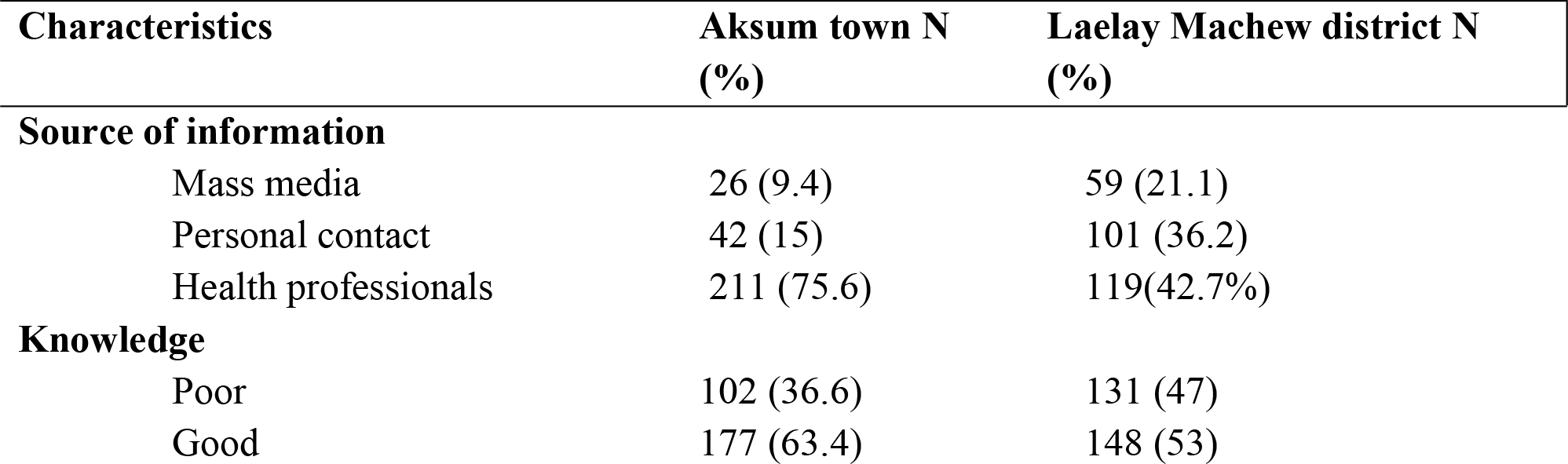
Personal and health service characteristics of dog owners in Aksum town and Laelay Machew district, Tigray region, Ethiopia, March 01-20/2018

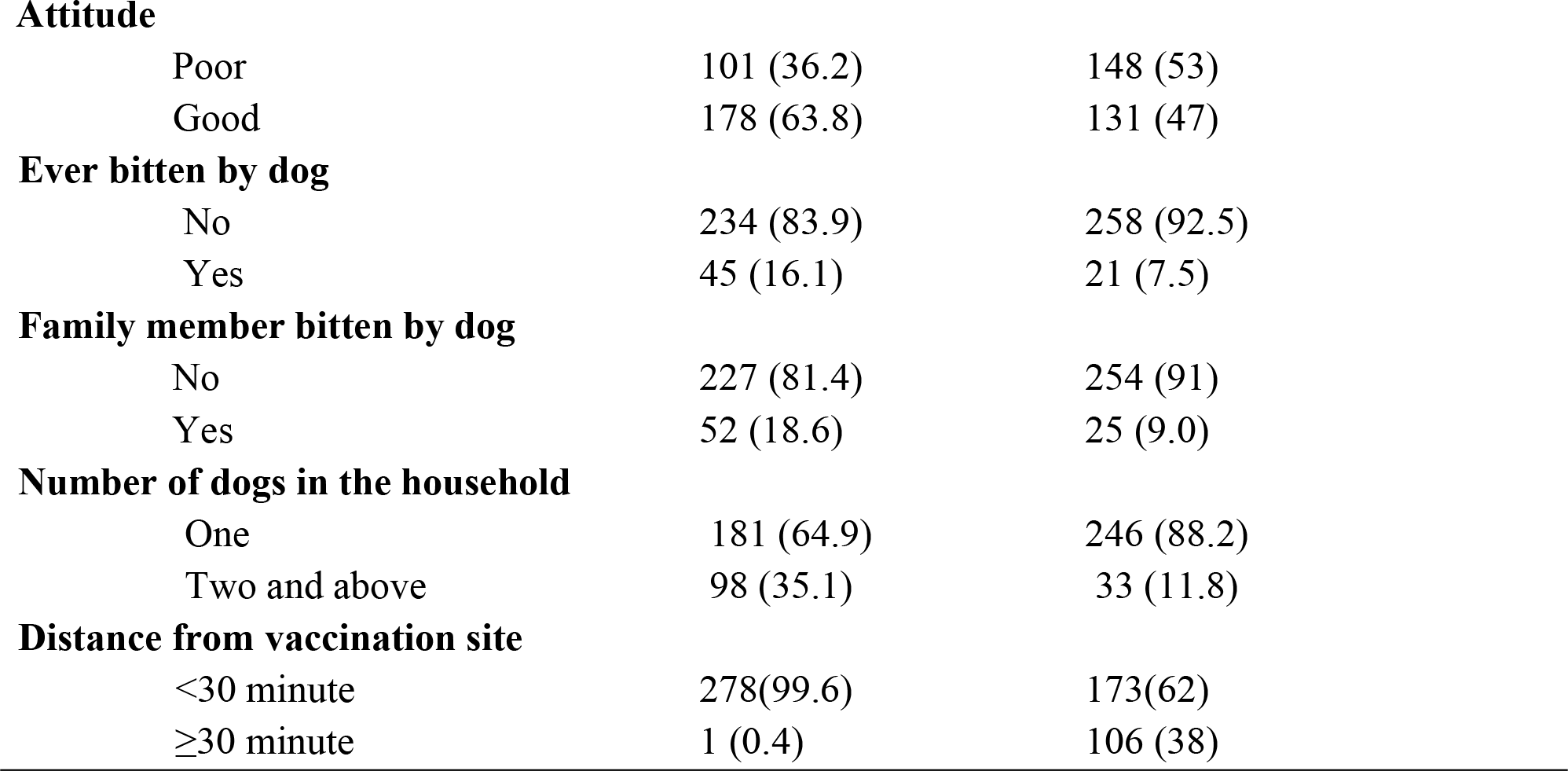

Nineteen (6.8%) of urban respondents and nine (3.2%) of rural respondents would wash the dog bite wound with water and soap. About 43 (15.4%) in urban and 37 (13.3%) in rural respondents prefer traditional healers. Of the total dog owners, 167 (60%) of urban and 88 (31.5%) of rural respondents had ever vaccinated their dogs and presented dog vaccination certificates. But 55% of both urban and rural respondents vaccinated their dogs within the past 12 months. About 85(30.5%) of urban respondents and only eight (3%) of rural respondents replied that their dogs were free roaming (Table 4).

**Table 4:** Rabies prevention and control practice among dog owners in Aksum town and Laelay Machew district, Tigray region, Ethiopia, March 01-20/2018

| Practice questions | Aksum town N (%) | Laelay Machew district N (%) |
| --- | --- | --- |
| **Action for bite wound as first aid** |  |  |
| Wash with water & soap & apply alcohol | 19 (6.8) | 9(3.2) |
| Visit health facility | 206 (73.8) | 229 (82.1) |
| Inform local leaders | 10 (3.5) | 4(1.4) |
| Go to traditional healers | 43 (15.4) | 37 (13.3) |
| Do nothing | 1 (0.3) | 11(3.9) |
| **Timing of hospital presentation** |  |  |
| Immediately | 227 (81.4) | 211(75.6) |

|  |  |  |  |
| --- | --- | --- | --- |
|  | 02-14 day | 24 (8.6) | 36 (12.9) |
|  | >14 day | 27(9.7) | 32 (11.5) |
| Is your dog vaccinated? |  |  |  |
|  | No | 112(40.1) | 191(68.5) |
|  | Yes | 167(59.9) | 88 (31.5) |
| **Date of vaccination** (checked in certificate) |  |  |  |
|  | <12 month | 92 (55) | 49 (55.7) |
|  | ≥12 month | 75 (45) | 39 (44.3) |
| **Action taken for suspected rabid dog** |  |  |  |
|  | Report to Vet. Officers | 106 (38) | 119 (42.7) |
|  | Killing | 122(43.7) | 151(54.1) |
|  | Do nothing | 51(18.3) | 8 (2.9) |
| **Action to rabid animal carcasses** |  |  |  |
|  | Throw it away | 9 (3.2) | 7 (2.5) |
|  | Bury/burn | 145(52) | 150 (53.8) |
|  | Do nothing | 19 (6.8) | 10 (3.6) |
| **How do you restrain your dog** (observe) |  |  |  |
|  | Tied whole day | 124 (44.4) | 210 (75.3) |
|  | Tied half day | 70 (25.1) | 60 (21.5) |
|  | Free roaming | 85(30.5) | 8 (2.9) |
| **Overall** | **Poor (<mean score)** | **156(56)** | **174(62.4)** |
| **Practice** | **Good (≥ mean score)** | **123(44)** | **105(37.6)** |

### Factors associated with rabies prevention and control practice

In the bivariate analysis; variables such as age, marital status, knowledge, source of information, occupation, household size and attitude of respondents were found to be associated with poor rabies prevention and control practice in Aksum town (Urban). Variables such as sex, age, marital status, educational level, number of dogs owned, distance from dog vaccination site, source of information, attitude, his/her history of dog bite and any family bitten were found to be associated with poor practice in Laelay Machew district (Rural) at p-value ≤0.25.

Two different models were fitted to identify factors associated with poor rabies prevention and control practice among urban and rural owners. The first model was fitted only for urban dog owners. Accordingly statistically significant association was observed having poor practice with occupation and attitude of respondents. Being government employees AOR (95% CI) = 0.35(0.13, 0.94) and private employees AOR (95% CI) = 0.39 (0.16, 0.97) were 65% and 61% less likely to have poor practice respectively than farmers. Respondents having poor attitude AOR (95% CI) = 1.84 (1.04, 3.25) were two times more likely have to poor practice than counterparts (Table 5).

**Table 5:**
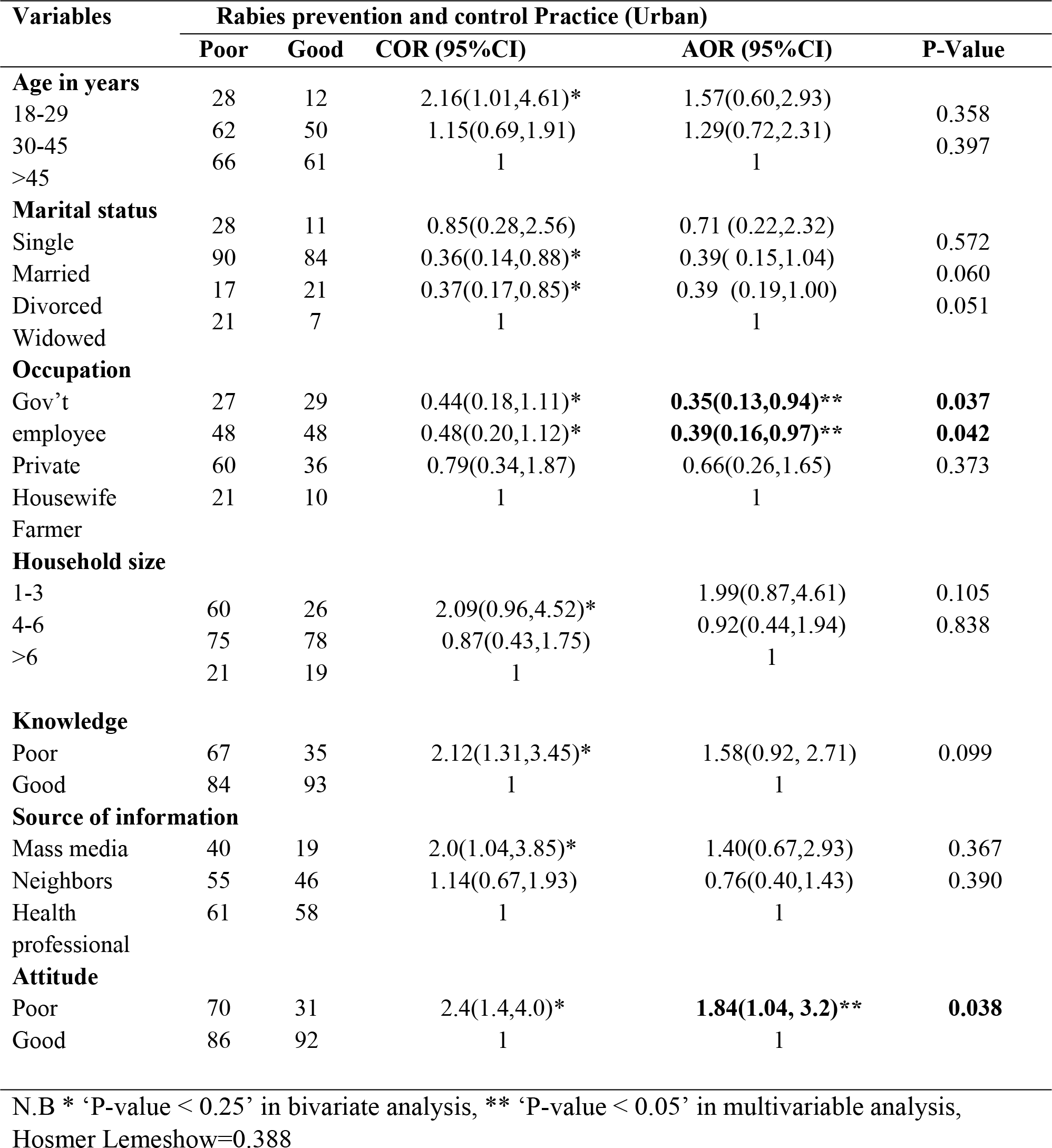
Factors associated with poor rabies prevention and control practice among dog owners in Aksum town, Tigray region, Ethiopia, March 01-20/2018

The second model was fitted for rural dog owners. Variables such as level of education, attitude, number of dogs, distance from vaccination site, dog bite history of family members and source of information were significantly associated factors with the outcome variable.

Respondents having no formal education AOR (95% CI) =6.41 (1.1, 38.6) and primary school AOR (95% CI) =10.4 (1.7, 61.6) were 6.4 times and 10.4 times more likely to have poor practice than counterparts. Those who have poor attitude AOR (95% CI) =2.19 (1.18, 4.05) were two times more likely to have poor practice than counterparts. Households having only one dog AOR (95% CI) =3.31 (1.34, 8.15) were 3.3 times more likely to have poor practice than having two and above. Those travel ≥30 minutes to get dog vaccine AOR (95% CI) =4.26 (2.14, 8.47) were 4.3 times more likely to have poor practice than counterparts. Respondents having no history of dog bite among family member AOR (95% CI) =4.16 (1.49, 11.6) were 4.2 times more likely to have poor practice than counterparts. Those with relative or neighbors as source of information AOR (95% CI) =3.64 (1.31, 10.1) were 3.6 times more likely to have poor practice than health professionals (Table 6).

**Table 6:**
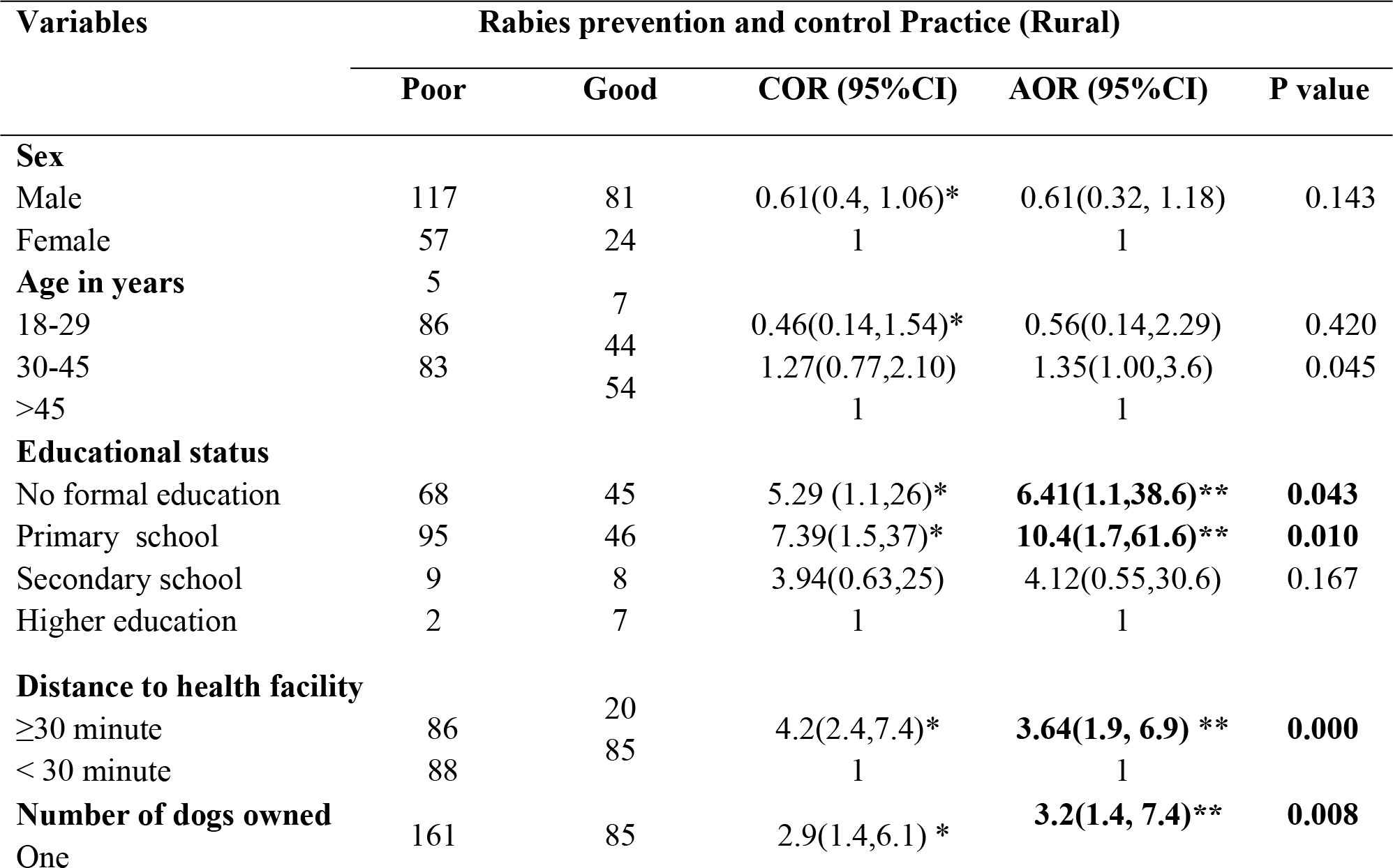
Factors associated with poor rabies prevention and control practice among dog owners in Laelay Machew district, Tigray region, March 01-20/2018

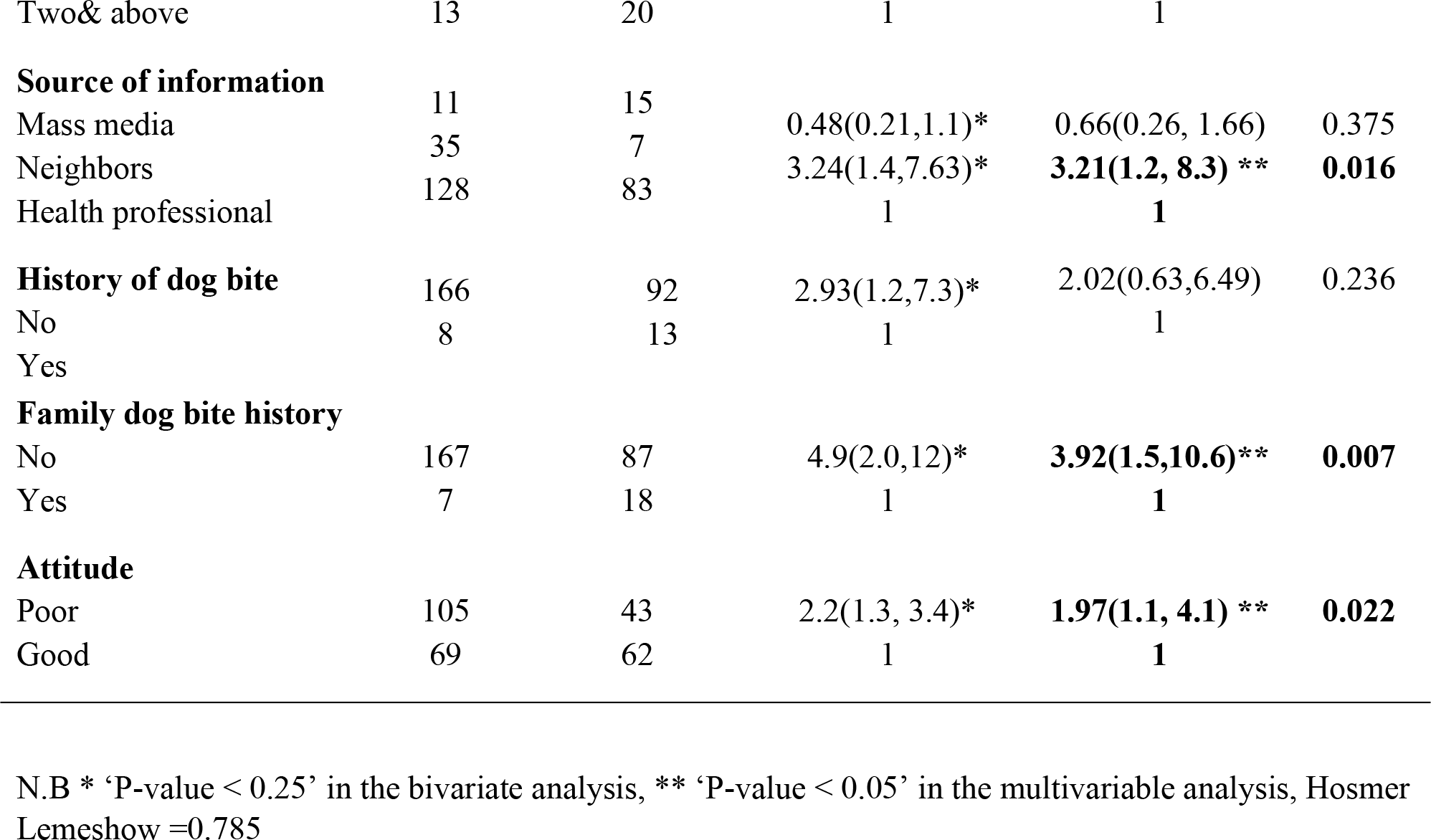

The third model was fitted to identify the overall factors associated with poor rabies prevention and control practice in both urban and rural communities. According to bivariate analysis; variables such as sex, marital status, occupation, educational status, household size, number of dogs owned, source of information, distance from vaccination site, dog bite history of respondents and family members, attitude, and place of residence were found factors as associated rabies prevention and control practice of dog owners at P value ≤0.25.

In multivariable analysis; occupation, distance, attitude and place of residence were significantly associated with the outcome variable regardless of the place of residence. Respondents who were government employee AOR (95% CI) = 0.40 (0.18, 0.89) and private employee AOR (95% CI) =0.34 (0.20, 0.70) were 60% and 66% less likely to have poor practice than farmers. Respondents travel ≥30 minute to get dog vaccine AOR (95% CI) =3.38 (1.90, 6.02) were 3.4 times more likely to have poor practice than counterparts. Respondents having poor attitude AOR (95% CI) =2.25 (1.54, 3.28) were 2.2 times more likely to have poor practice than counterparts.

**N.B:** Being rural resident AOR (95% CI) = 2.31 (1.18, 4.51) was 2.3 times more likely to have poor practice than urban ones.

## DISCUSSION

The finding of this study showed that the overall prevalence of poor rabies prevention and control practice among dog owners was 56% in Aksum town (urban) and 62.4% in Laelay Machew district (rural) with statistical difference in prevalence among urban and rural dwellers. This is higher than the study done in and around Ambo town in 2017 (36.5%) [29]. This might be due presence of poor community attitude towards rabies prevention and control in the current study (53%) which was 49.5% in Ambo town. Moreover; the poor practice was as a result of lower dog vaccination coverage (55%) than WHO recommendation of annual rabies vaccination coverage for herd immunity at least 70% [2].

Among urban dog owners; statistically significant association was observed in level of practice with occupation and attitude of respondents. Being government and private employees were less likely to have poor practice on rabies prevention and control than farmers. This is consistent with the study done in Nigeria and Debretabor town [19, 30]. This might be probably due to the government employee have higher educational background enable to access health information than farmers. Respondents having poor attitude were more likely to have poor practice than counterparts that is consistent with the study conducted in India [31]. This implies positive attitude drives to practice well.

Whereas among rural dog owners; households having only one dog were more likely to have poor practice than two and above. This might be to prevent cross infection of two and above dogs. In other way might be due to one vaccine vial contains five doses opened for five dogs come at the same time but most time it is difficult for those having only one dog. This implies less attention was given to one dog than two and above.

Respondents who travel ≥30 minute to get dog vaccination site were more likely to have poor practice than the counterparts. This might be due to not easily access to health education programs, vaccination schedules and timing of hospital presentation. Respondents who did not have any family exposure to dog bite were higher to have poor practice than having exposure history. This is consistent with the study done in Kakamega, Kenya and Hawasa [17, 18, 24]. This implies having exposure history helps to have good practice towards rabies prevention and control. But everybody need to aware of prevention both before and after bite exposure.

Respondents with no formal education and attended primary schools were more likely to have poor practice than counterparts. This was consistent with the KAP studies conducted in Dedo district [32], Addis Ababa[33], Debretabor town [19], Hawasa town [24], and Debark district[20]. The possible explanation could be educated persons would have better information access and can easily understand the disease prevention and control measures. Schools have great role in acquisition of comprehensive knowledge on the disease.

Respondents that their sources of information got from personal contacts (relatives or neighbors) were 3.2 times more likely to have poor practice than heard from health professionals. This may be due to lack of appropriate and comprehensive information on rabies transferred from relatives/neighbors. Respondents having poor attitude were two times more likely to have poor practice than counterparts.

Generally; rural dog owners were two times more likely to have poor rabies prevention and control than urban ones. This is consistent with the study in Tanzania [16, 34]. This might be due to behavioral change of urban dog owners on early hospital presentation of dog bite victims since they were living nearby to health facility and had better access to health information from different sources.

### Strength and limitation of the study

As strength of the study; this is community based comparative cross-sectional study design of urban and rural communities with high (100%) response rate, however; there could be recall bias of respondents. To minimize the recall bias we included observation part to cross check the dog vaccination certificate and tying of dogs during time of data collection.

## CONCLUSION

This study revealed that poor rabies prevention and control practice was higher among dog owners in Laelay Machew district than Aksum town. Occupation and attitude of dog owner households were identified as independent associated factors for rabies prevention and control practice in Aksum town. While Level of education, distance from dog vaccination site, number of dogs in the household, history of dog bite any of family member, source of information and attitude of respondents were identified in Laelay Machew district.

Then, it need to expand and strengthen the outreach dog vaccination campaigns especially to the households travel ≥30 minutes to get vaccination to increase vaccination coverage and all dog owners with dogs aged three months and above should be encouraged to take their dogs for rabies vaccination. And collaboration is basic with veterinary office and other sectors need to strengthen and expand outreach health education on rabies prevention activities.

**Figure 1:**
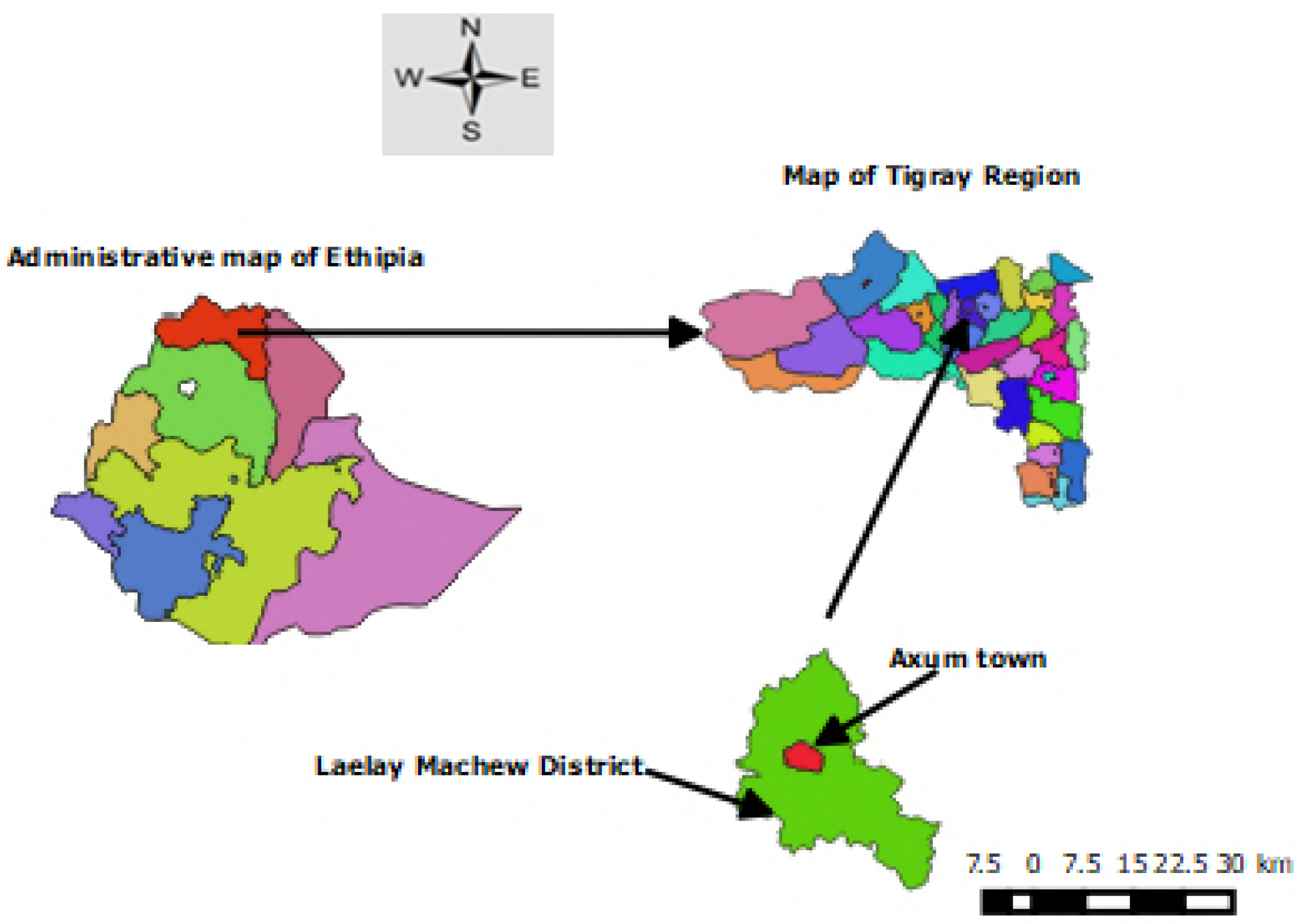
Administrative map of Aksum town and Laelay Machew district, Tigray, Ethiopia, 2018.

**Figure 2:**
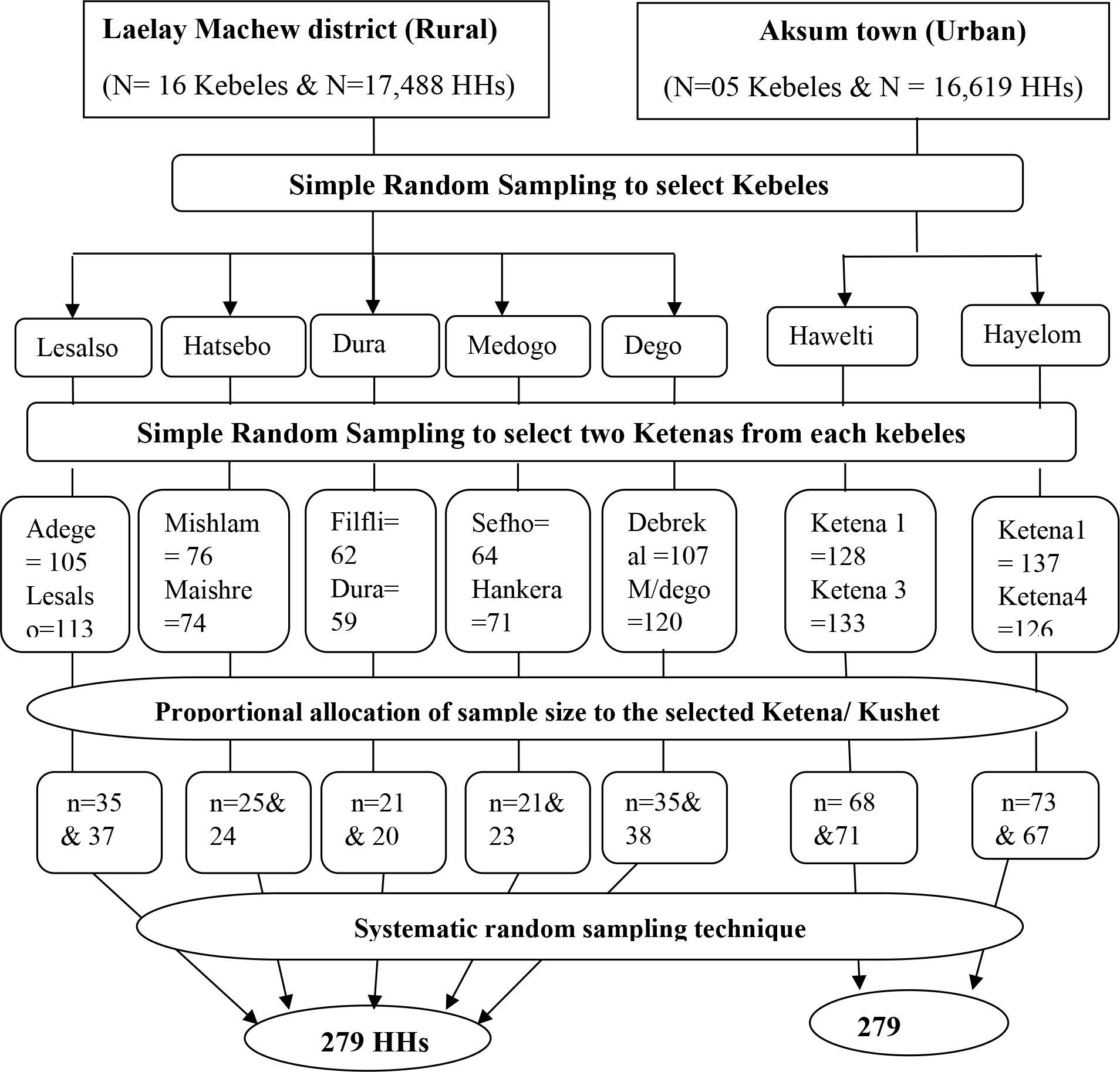
Schematic diagram of sampling procedure for selection dog owner households in Aksum town and Laelay Machew district, Tigray, Ethiopia.

## Declarations

### Ethics approval and consent to participate

Ethical clearance was obtained from ethical review board of Mekelle University, College of Health Sciences. Support letter was obtained from Tigray Regional Health Bureau, Axum town and Laelay Machew district agriculture and health offices. Informed verbal and written consent was obtained from study participants after detailed explanation on the purpose, possible risks and benefits of the study before data collection. Participants also informed the right to withdraw from interview during data collection. Confidentiality of information was secured.

### Consent for publication

Not Applicable.

### Availability of data and materials

All the data supporting the findings is contained within the manuscript, when there is in need the data-set used for the present study’s conclusion can be accessible from the corresponding author on reasonable request.

## Authors’ contributions

LG is the principal investigator of this study who conceptualized the study and with AGB, GB and AA he contributed; recruited study participants, funding acquisition and made data collection, providing methodology, study investigation and supervision, data analysis, validation and writing (original draft preparation review and editing), preparing and review of manuscript.

## Acknowledgment

We would like to thank to Mekelle University and Tigray regional health bureau, and Ethiopian ministry of health. We have a grateful thanks to Aksum and Laelay Maychew health offices staffs for giving me needed data and material. We have special thanks to my data collectors and study participants.

